# SNP Data Quality Control in a National Beef and Dairy Cattle System and Highly Accurate SNP Based Parentage Verification and Identification

**DOI:** 10.1101/148171

**Authors:** M.C. McClure, J. McCarthy, P. Flynn, J. McClure, K. O’Connell, J.F. Kearney

## Abstract

A major use of genetic data is parentage verification and identification as inaccurate pedigrees negatively affect genetic gain. Since 2012 the international standard for single nucleotide polymorphism (SNP) based verification in *Bos taurus* cattle has been the ISAG 100 and 200 SNP panels. While these SNP sets have provided an increased level of parentage accuracy over microsatellite markers (MS), they can validate the wrong parent for an animal at ≤1% misconcordance rate levels, indicating that more SNP are needed if a more accurate pedigree is required. With rapidly increasing numbers of cattle being genotyped in Ireland that represent 61 *Bos taurus* breeds from a wide range of farm types: beef/dairy, AI/pedigree/commercial, purebred/crossbred, and large to small herd size the Irish Cattle Breeding Federation (ICBF) analysed different SNP densities to determine that at a minimum ≥500 SNP are needed to consistently predict only one set of parents at a ≤1% misconcordance rate. For parentage validation and prediction ICBF uses 800 SNP selected based on SNP clustering quality, ISAG200 inclusion, call rate (CR), and minor allele frequency (MAF) in the Irish cattle population. Large datasets require sample and SNP quality control (QC). Most publications only deal with SNP QC via CR, MAF, parent-progeny conflicts, and Hardy-Weinberg deviation, but not sample QC. We report here a genomic sample QC pipeline to deal with the unique challenges of >1,000,000 genotypes from a national herd such as SNP genotype errors from mis-tagging of animals, lab errors, farm errors, and multiple other issues that can arise.

## Introduction

Since the 1960’s bovine pedigree verification has been performed with various DNA technology, initially performed with blood groups (Stormont, 1967), then microsatellite markers (MS) (Davis and DeNise, 1998), and now transitioning to single nucleotide polymorphisms (SNP). While the initial cost and availability of each new technology has hindered their adaption, their increasing ability to reduce pedigree errors cannot be ignored. A 10% pedigree error rate can have a 6-13% effect on the inbreeding coefficient, 11-18% reduction on breeding value trends, 2-3% loss in selection response (Visscher et al., 2002a, Banos et al., 2001), and a downward basis on heritability estimates (Israel and Weller, 2000). While sire error rates have been estimated at >7% in national herds, dam errors and missing parental information can be substantial, especially in commercial herds (Sanders et al., 2006, Harder et al., 2005) and their effects are additive (Sanders et al., 2006).

With all technology there is a need to balance cost with performance, for parentage validation, the question has typically been how many markers are needed to obtain a high probability, but not necessarily 100%, that the reported parentage is correct. The International Society of Animal Genetic (ISAG) recommended parentage SNP panel of 100 SNP (ISAG100) has a reported parental exclusion probability (PE) of >0.999 and the ISAG200 panel (200 SNP) has a PE >0.9999999 (http://www.isag.us/). Many groups world-wide primarily or only use the ISAG100 or ISAG200 panel for initial bovine parentage validation, some use less. While the PE values for the ISAG SNP panels appear sufficient, Vandeputte (Vandeputte, 2012) notes that many reported PE values are overly optimistic. Increasing numbers of markers are needed to maintain the same PE value as the population size increases, and a marker set with a high PE value can still have a low probability of complete exclusion of all false parentage with large populations [9].

These issues with PE values could be one of the reasons why we and others have reported (McClure et al., 2015, Strucken et al., 2015) that using lower density parentage SNP panels like the ISAG100 and ISAG200 can result in false-positive validations and result in multiple parents being predicted when used in large population datasets. While there currently is no international standard for which or how many SNP to use for parentage outside of the ISAG set, we argue that a larger SNP set should be used such as the 800 SNP that ICBF uses for parentage validation and prediction was selected, and has proved, to be highly accurate and useful for sample quality control (QC) and parentage across multiple *Bos taurus* breeds.

As ICBF’s genotype database has rapidly increased since 2013, from ~25,000 to currently having >1,000,000 animals genotyped, data quality has become a higher concern, especially as ‘once in a million’ and ‘rare’ types of errors are encountered. Most publications only perform SNP and individual genotype QC based on SNP QC via call rate (CR), minor allele frequency (MAF), and Hardy-Weinberg deviation (Turner et al., 2011). While Wiggans et al. (Wiggans et al., 2011) has previously briefly describe additional QC steps they used, ICBF has developed their own QC pipeline to deal with other issues, some foreseen others unique, which we describe in full below. We hope that the full description of our sample QC pipeline and parentage process will be useful to other groups as they grapple with larger genotyping datasets, regardless of the species. While most errors come about due to accidents or non-malicious actions, but a very small amount could be due to intentional actions.

The main QC concerns for ICBF are 1) is this genotype ok and 2) does the genotype really belong to the listed animal. While most errors come about due to accidents or non-malicious actions, a very small amount could be due to intentional actions. We have tried to design a QC pipeline will identify both types of errors once enough data is collected. An example of potential errors includes:

A. Farmer

1. Calmer animal sampled as requested animal is dangerous
2. Same animal sampled >1 times but labelled differently
3. Wrong animal sampled
B. Laboratory

1. Genotype duplicate due to technician error on sample
2. Genotype assigned to wrong sample
C. AI centre

1. Wrong label attached to AI straw
2. Wrong animal sampled
D. Genotype format

1. Type (AA, AB, or BB) is missing, or low frequency
2. Format is wrong, e.g. mix of AB and ACTG format

## Material and Methods

Animal Care and Use Committee approval was not obtained for this study because the data were obtained from the existing Irish Cattle Breeding Federation (ICBF) database, Bandon, Co. Cork, Ireland

### ICBF animal and genotype database

The ICBF database was set up in 1998 and holds numerous records on all dairy and beef cattle in Ireland including, but not limited to, date of birth, sex, reported dam, reported sire or sire breed, parentage validation status and method, animal movement, date of death, pedigree based breed composition, milk recordings, carcass data, along with multiple other phenotypes. Eighty-five *Bos taurus* breeds are represented in Ireland, 58 beef and 27 dairy, with 44 breeds having at least 1 purebred animals genotyped in the ICBF database and 61 breeds being represented in a purebred or crossbred genotyped animal (Table S1). Data is reported to the database from, but not limited to producers, Department of Agriculture, Food, and Marina (DAFM), marts, abattoirs, veterinarians, AI technicians, milk co-ops, and herd books. SNP genotype data is also reported to ICBF on Irish animals from commercial genotyping labs and via international collaborations. At submission the ICBF database holds valid genotypes on >1,000,000 individuals, while animals with pedigree and phenotype records stands at >40.5 million.

Genotyped animals represent a mixture of male and female Irish beef, dairy, purebred, crossbred, pedigree, and commercial animals from 59 *Bos taurus* breeds who were genotyped on multiple SNP chips such as the Illumina 3K, LD, 50K, and HD; GeneSeek GGPLD and HD; and our custom International Beef and Dairy (IDB) chip (Mullen et al., 2013, Illumina Inc, 2011a, Illumina Inc, 2009, Illumina Inc, 2010, Matukumalli et al., 2009, Neogen Corporation, 2012, Neogen Corporation, 2013, Illumina Inc, 2011b). As the Illumina 3K has not been commercially available since September, 2011 (personal communication, André Eggen, 23/2/2015), and with its higher variability in genotyping accuracy (Wiggans et al., 2012) 3K genotypes were not used for the analysis or pipeline listed below

### Parentage: SNP Panel and Process

#### Minimum SNP needed to generate 1 predicted sire

This work was originally described by us (McClure et al., 2015), and is briefly described here. In March, 2014 an analysis was done to determine the minimum number of SNP needed to predict only 1 sire using 65,456 animals that had been genotyped across multiple commercial Illumina SNP chips including the IDB (v1 and v2). MAF on the 6,912 SNP from the Illumina LD base content that are present across the above mentioned chips, were calculated across all genotyped animals in the ICBF database (Table S2). Panels of SNP (200, 300, 400, 500, 600, 700, 800, 900, 1000, 1100, and 1250) were designed that contained the ISAG200 SNP and increasing numbers of top MAF ranked SNP.

The number of predicted sires at the 0.5-1% misconcordance rate for each SNP panel for 7,092 animals with SNP validated sires, was determined via running them through an in-house designed parentage prediction program with the 65,456 reference animals. The prediction program returned all sires and dams that had less than the set number of SNP misconcordances (≤1.0% for ≤500 SNP, ≤0.5% for ≥600 SNP), and were >18 months older than the animal. The breed, pedigree status, herd, farm location, and AI status of the animal and predicted parents were not considered, only age difference to the animal, predicted animal’s sex, and misconcordance rate was considered.

#### Parentage SNP Revaluation 1

This work was also originally described by us (McClure et al., 2015), and is briefly described here. In March, 2015 >180,000 genotyped animals were in the ICBF database, SNP CR and MAF for the core LD SNP were analysed to determine if any SNP should be dropped or added to the original 2014 800 Parentage SNP set (ICBF800v1). High rates of misconcordance errors in IDBv2 genotype data from ISAG parentage SNP ARS-USMARC-PARENT-DQ837645-RS29015870 directed us to analyse SNP cluster patterns for all ISAG200 SNP and the top 1,000 MAF SNP. Cluster issues were analysed to see if they were breed specific or independent and if they were present on other Illumina SNP chip data. An updated set of 800 parentage SNP (ICBF800), very similar to the previous set, and SNP ranking by ISAG listing and Irish MAF was created and used in the remaining work.

#### Parentage SNP Revaluation 2

The difference accuracy levels using the ICBF800 SNP or fewer SNPs were evaluated in August 2016 with >570,000 genotypes available in the ICBF database. 300,020 animals whose listed sire or dam was SNP genotyped were used; 19,774 had both parents SNP genotyped. Subsets of the ICBF SNP set: 100, 200, 300, 400, 500, 600, 700, and 800 SNP, were developed by using the ISAG 100 and ISAG200 sets and then adding additional SNP in descending order of their March 2015 MAF. Any SNP listed in Table S2 with quality control (QC) issues were not used. SNP misconcordances between the animal and the original listed or predicted parent were recorded for each SNP density.

#### Parentage SNP Revaluation 3

In December, 2016 we also addressed if animals in the ICBF800 misconcordance result ‘grey zone’ of 5 to 12 mismatches, which represent 0.5% to 1.5% misconcordance levels, should be further analysed. While this was not a concern in the past, as ICBF approached 1 million genotyped animals a very small percent of animals have appeared in this grey zone of doubtful parentage result. In December 2016 845,275 genotyped animals were in the ICBF database and the number of SNP misconcordances with their original listed parents were used to analyse the frequency of each SNP misconcordances level, ‘grey zone’ counts, and what should be done about ‘grey zone’ results.

#### Parentage Validation, Prediction, and Suggestion Pipeline

All animals with valid SNP genotypes received by ICBF that pass the genotype QC process, listed below, are automatically sent through the Parentage Pipeline. Depending on the animal type (commercial, pedigree) and owner or herd book requests an animal will exit the pipeline at different points. SNP based parentage validation and prediction are performed using the ICBF800 SNP described above. Microsatellite imputation (Mcclure et al., 2013) is performed using a set of 908 SNP for pedigree animals whose parentage is not SNP validated or predicted. When requested to suggest potential non-genotyped parents a Genetic Relationship Matrix (GRM) via SVS software (Golden Helix, Montana, USA) is performed using the Illumina LD SNP panel to identify close genetic relatives. A GRM analysis will be performed on either all genotyped animals from a farm or all purebred animals from a select breed as due to current software limitations SVS can only perform a GRM on ~70,000 animals for ~6,900 SNP.

## Genotype QC Pipeline

Genotype quality is one of the easiest and most used QC tools used. At ICBF we use a standard animal call rate (CR) of ≥0.90 for a genotype to be used. The calculation of this individual CR does not include SNP that have CR <0.85 across our database. While genotypes received should all be in a standard format, for instance Illumina AB allele or Top ACTG allele format, ICBF has received genotypes before that have had a mix of genotype formats or missing genotypes, such as no BB genotype. The percent of each genotype an animal has can be used to access genotype quality as well. To determine acceptable frequencies of each genotype 846,868 animals with call rates ≥0.90 from multiple chip types were analysed. A genotype must pass all genotype QC quality checks to be used downstream.

## Animal QC Pipeline

### SNP Duplicate Check

Excluding identical twins, two animals should not have the same genotype across large numbers of SNP. When they do this is an indication of either a farm or lab level error. The former being that the same animal was sampled twice, possibly with a decent amount of time between tissue collections when a different animal was requested, the latter usually occurring within a narrow time frame. Lab errors can often be detected by analysing the genotypes of all animals processed during a set time frame, such as daily or weekly. Farm error identification requires analysing all of the farm’s DNA genotypes. Finally you could have a farm level error combined with a mis-labeling of the DNA collection device (usually labelled by a 3^rd^ party), to detect these you would need to analyse all genotypes in the national database.

Comparing the full SNP genotype of an animal against a large number of records is possible, but far too time consuming. To speed up this process, we use the ICBF800 SNP set to identify potential SNP duplicates and then compare all available SNP to confirm which potential duplicates are true duplicates. Other QC information could allow for one to determine who the genotype truly belongs to.

### Animal with >1 non matching genotypes

Animals can be genotyped more than once within and across countries for multiple legitimate reasons. Regardless of the number of times an animal is genotyped any SNP common across the SNP platforms and chip types should generate the same genotype, for instance Illumina and Affymetrix SNP chips have >99% concordance rates across platform and tissue types (McClure et al., 2009, Montgomery et al., 2005, Feigelson et al., 2007, Woo et al., 2007). If the concordance rate between two genotypes assigned to an animal is <99% then this indicates they are probably not from the same animal and one should be able to determine which genotype truly belongs to the individual. The ICBF800 panel is used to identify potential issues and then all available genotypes are used to confirm. Other QC information could allow for one to determine which genotype correctly belongs to the animal.

### Sex Prediction and Pseudoautomal Region of Chromosome X Determination

Sex prediction can be performed using chromosome Y (chrY) or chromosome X (chrX) SNP that are located in the non-pseudoautosomal (nPAR) region. Sex prediction using only chrY SNP is logically simpler to use, but not all commercial SNP chips contain chrY SNP. Sex prediction only using the heterozygosity rates of chrX SNP can be more challenging as care must be taken to avoid using PAR SNP and highly inbred females such, as L1 Dominette 01449 (Henderson et al., 2005), can look like males if not enough SNP are used.

As the Illumina LD base content is widely used across multiple commercial and custom bovine SNP chips the 9 chrY SNP from the LD chip were selected inclusion on the IDBv1 for chrY sex prediction: BOVINEHD3100000048, BOVINEHD3100000099, BOVINEHD3100000103, BOVINEHD3100000210, BOVINEHD3100000515, BOVINEHD3100000517, BOVINEHD3100001188, BOVINEHD3100001404, and BOVINEHD3100001406. By 2014 ICBF had 4,901 HD genotyped animals, 232 of them being female and 28,317 IDBv1 genotyped animals, 3141 being female. Of those 9 chrY SNP from the LD base, BOVINEHD3100000515 and BOVINEHD3100001404 were heterozygous in 280 and 503 HD genotyped and 469 and 1090 IDBv3 genotyped males, respectively. None of the 9 chrY SNP were present in the genotyped females, excluding 107 IDBv1 females for BOVINEHD3100000515. Therefore only the remaining 7 chrY SNP were used for sex prediction.

SNP in the PAR of the X chromosome was determined by analysing chrX allele heterozygosity rates in 37,281 males and 568,841 females who were genotyped on the IDBv3 and had 7 or 0 called chrY genotypes, respectively, and had ≥90% CR for 491 chrX SNP. After filtering for unique location on chrX (UMD3.1 assembly) 467 SNP remained. MAF and CR were calculated for these SNP across the 606,122 animals and any SNP with MAF <0.01 or CR < 0.90 was excluded, leaving 391 SNP. The PAR region was determined via SNP with high male heterozygosity rates and genomic positions. SNP MAF rates were determined for the nPAR and PAR in both males and females. Individual heterozygosity rates were determined for males and females for PAR and nPAR using SVS (Golden Helix).

### Breed Composition Prediction

The breed composition of an animal is predicted via Admixture (Alexander et al., 2009) using 36,819 SNP that map to autosomes on the UMD3.1 assembly (Zimin et al., 2009)and are common across the 50k, HD, and IDBv3 panels. The reference population of is comprised of 22,610 purebred animals from fourteen breeds, Angus, Aubrac, Blonde D’Aquitane, Belgian Blue, Charolais, Fresian, Hereford, Holstein, Jersey, Limousin, Parthenaise, Salers, Shorthorn, and Simmental, with a minimum of 600 and a maximum of 2,000 animals per breed.

Animals genotyped on lower density chips were not used in the reference population to maximize the number of SNP used because their inclusion would not have increased the number of reference breeds, nor greatly increased the animals in breeds with <2,000 animals. Breeds with >2,000 genotyped purebreds had their breed composition reference animals selected randomly. Breeds with lower numbers of purebred genotyped animals were tried but not used do to their low prediction accuracy, often <50% correlation to the animal’s reported breed composition.

### Parentage

National bovine pedigree errors have been reported between 4-13% (Visscher et al., 2002b, Leroy et al., 2012) based upon farmer recorded information, in Ireland, a 7-9% listed pedigree error rate is seen on a national level. While one listed parent may be listed incorrectly, it is rare that both listed parents would be wrong. Even less likely, though possible, is that the true parent(s) do not reside on or close to the farm, excluding AI sires. While the true sire might be an intact young bull, or neighbouring stock bull the sire is usually geographically located in the same area, excluding AI sires. ICBF takes into account the geographic location of the sire and dam, along with their breeds, to determine if the predicted parents for an animal are logical.

In addition to individual parentage validation and prediction checks, if both parents SNP validate one can also check to see if the mating could have occurred from them. This is done by checking that for each parentage SNP the calf is homozygous for that both parents are not homozygous for the same allele. So if the calf is AB for a SNP then the sire and dam both cannot be AA or BB. If >0.5% of the ICBF800 parentage SNP have this type of misconcordance the parentage is checked using all possible SNP and a flag is raised for manual checking.

As a genotype can appear correct, having passed SNP QC and the already listed Animal QC steps, it can still be wrongly assigned to an animal. For instance, if fraternal twins, of the same sex, are genotyped but the genotype is assigned to the other animal. In these cases the error will only become apparent when their offspring are genotyped.

### Additional Flag

While possible if a tissue sample is sent in to be genotyped >45 days after the animal’s death is recorded ICBF records this as a minor warning flag. The genotype is not invalidated by this but this flag is taken into account for genotype resolution if needed, for instance for SNP duplicates. AI sires are excluded from this flag if an AI straw is the submitted sample.

## Results

### SNP allele rates

For all genotype classes <0.0001% of animals had a genotype (AA, AB, or BB) frequency lower than 20%. Out of all 846,868 animals analysed only 76 had a genotype class with a frequency ≤20%, and only 5 had a frequency ≤15% (Table 1). As far as we know this is the first time the pattern of individual genotype frequencies has been analysed. While highly inbred animals would have reduced AB frequency, this analysis strongly indicates any animal with a genotype class frequency <20% should be flagged for further analysis.

**Table 1.** Count of animal’s genotype frequency for AA, AB, and BB genotypes from 846,868 genotyped individuals with a genotype CR>0.9 and all 3 genotypes present.

| Frequency | AA | AB | BB |
| --- | --- | --- | --- |
| 0 | - | - | - |
| 0.005 | - | - | - |
| 0.01 | - | - | - |
| 0.02 | - | - | - |
| 0.03 | 1 | - | - |
| 0.04 | - | - | - |
| 0.05 | - | - | - |
| 0.06 | 1 | - | - |
| 0.07 | - | - | - |
| 0.08 | 2 | - | - |
| 0.09 | - | - | - |
| 0.1 | - | - | - |
| 0.15 | - | 1 | - |
| 0.2 | 2 | 68 | 1 |
| 0.25 | 1977 | 2745 | 6 |
| 0.3 | 620489 | 57372 | 44438 |
| 0.35 | 223499 | 564809 | 153114 |
| 0.4 | 885 | 78550 | 637289 |
| 0.45 | 2 | 138927 | 12002 |
| 0.5 | - | 4385 | 9 |
| >0.5 | 1 | 11 | 9 |

Genotypes with missing genotype classes, often BB, have been observed when genotypes were obtained from other scientific groups. These genotypes are invalidated and not included in the genotype class frequency analysis.

### Parentage SNP Panel

MAF and CR analysis resulted in the identification of SNP present in the core Illumina LD panel that were highly informative (MAF >0.45) for parentage across multiple breeds in Ireland. Analysis of the Illumina BeadStudio files for the ISAG200 and top 1000 MAF SNP identified 15 SNP with clustering issues (Table S2, Figure S1) that were only apparent at high genotyping throughput rates of thousands of animals a week. The clustering issues were not breed or sex dependent. Five ISAG parentage SNP are noted in Table S2 that have either very low MAF (<0.001), very low CR (<0.3), or clustering issues. A retrospective analysis using data from other Illumina SNP chips (LD, 50k, HD, IDBv1-3) was performed on the ISAG200 SNP with clustering issues and similar cluster issues were seen when large numbers of animals were analysed (Figure S1).

Based on the March 2015 analysis, the ICBF800 SNP panel was designed for parentage validation and prediction was built using the ISAG200 SNP set as a base, minus the 5 listed in Table S2, and the next top 605 SNP based on MAF that did not have clustering issues. All animals in the ICBF database had their parentage validation reanalysed with the ICBF800 to provide a consistent parentage analysis for all animals. Since then the ICBF800 panel has been used by ICBF for parentage validation and prediction along with other QC checks.

The October 2016 analysis confirmed that using the ICBF800 results in greater accuracy than small SNP sets like the ISAG parentage SNP panel. If 100 (essentially the ISAG100 panel) had been used then 740 (0.23%) of the ICBF800 listed parentage results would have been different (Table 2). For 2.2 million animals, the average number of cattle born yearly in Ireland, this equates to ~10,000 parentage errors yearly if the ISAG100 panel was used instead of the ICBF800 panel. The yearly parentage error values would be higher if parentage predictions errors were included.

**Table 2.** Breed Composition reference population size and the correlation between the predicted and listed breed compositions from 713,814 animals Correlation on 710,000 animals between the predicted and listed breed composition for 14 breeds.

| Parent Validation<sup>1</sup> | AA<sup>2</sup> | AU | BA | BB | CH | FR | HE | HO | JE | LM | PT | SA | SH | SI | ave |
| --- | --- | --- | --- | --- | --- | --- | --- | --- | --- | --- | --- | --- | --- | --- | --- |
| NA | 0.9410 | 0.9493 | 0.9021 | 0.9036 | 0.9432 | 0.6220 | 0.9258 | 0.8799 | 0.9321 | 0.9385 | 0.8971 | 0.9497 | 0.9083 | 0.9272 | 0.9014 |
| Neither validated | 0.9044 | 0.9367 | 0.8752 | 0.7871 | 0.8869 | 0.4478 | 0.8907 | 0.7967 | 0.9219 | 0.8886 | 0.8607 | 0.9290 | 0.8775 | 0.8854 | 0.8492 |
| One validates | 0.9607 | 0.9520 | 0.9240 | 0.9435 | 0.9647 | 0.7306 | 0.9488 | 0.9135 | 0.9518 | 0.9617 | 0.9111 | 0.9606 | 0.9293 | 0.9492 | 0.9287 |
| Both validate | 0.9713 | 0.9656 | 0.9196 | 0.9385 | 0.9660 | 0.7265 | 0.9612 | 0.9408 | 0.7627 | 0.9642 | 0.9346 | 0.9706 | 0.9462 | 0.9583 | 0.9233 |

B) Animal's listed composition has some of the 14 breeds, N=711,579
| Parent Validation | AA | AU | BA | BB | CH | FR | HE | HO | JE | LM | PT | SA | SH | SI | ave |
| --- | --- | --- | --- | --- | --- | --- | --- | --- | --- | --- | --- | --- | --- | --- | --- |
| NA | 0.9417 | 0.9501 | 0.9035 | 0.9043 | 0.9432 | 0.6225 | 0.9261 | 0.8830 | 0.9344 | 0.9385 | 0.9096 | 0.9497 | 0.9113 | 0.9285 | 0.9033 |
| Neither validated | 0.9057 | 0.9381 | 0.8773 | 0.7894 | 0.8870 | 0.4483 | 0.8911 | 0.8032 | 0.9250 | 0.8885 | 0.8812 | 0.9291 | 0.8823 | 0.8878 | 0.8524 |
| One validates | 0.9610 | 0.9524 | 0.9250 | 0.9437 | 0.9648 | 0.7308 | 0.9490 | 0.9153 | 0.9529 | 0.9617 | 0.9185 | 0.9606 | 0.9308 | 0.9499 | 0.9297 |
| Both validate | 0.9717 | 0.9659 | 0.9205 | 0.9386 | 0.9660 | 0.7267 | 0.9614 | 0.9408 | 0.7720 | 0.9642 | 0.9411 | 0.9706 | 0.9474 | 0.9585 | 0.9247 |

C) Animal's listed composition is >50% of the 14 breeds, N=707,546
| Parent Validation | AA | AU | BA | BB | CH | FR | HE | HO | JE | LM | PT | SA | SH | SI | ave |
| --- | --- | --- | --- | --- | --- | --- | --- | --- | --- | --- | --- | --- | --- | --- | --- |
| NA | 0.9429 | 0.9511 | 0.9049 | 0.9049 | 0.9433 | 0.6245 | 0.9267 | 0.8836 | 0.9363 | 0.9386 | 0.9167 | 0.9499 | 0.9140 | 0.9303 | 0.9049 |
| Neither validated | 0.9075 | 0.9398 | 0.8792 | 0.7907 | 0.8872 | 0.4504 | 0.8922 | 0.8045 | 0.9273 | 0.8885 | 0.8921 | 0.9294 | 0.8864 | 0.8913 | 0.8547 |
| One validates | 0.9615 | 0.9531 | 0.9258 | 0.9440 | 0.9648 | 0.7318 | 0.9492 | 0.9155 | 0.9537 | 0.9617 | 0.9236 | 0.9607 | 0.9320 | 0.9508 | 0.9306 |
| Both validate | 0.9726 | 0.9662 | 0.9213 | 0.9389 | 0.9660 | 0.7291 | 0.9617 | 0.9410 | 0.7791 | 0.9643 | 0.9450 | 0.9707 | 0.9494 | 0.9590 | 0.9260 |

D) Animal's listed composition is only comprised from the 14 breeds, N=224,428
| Parent Validation | AA | AU | BA | BB | CH | FR | HE | HO | JE | LM | PT | SA | SH | SI | ave |
| --- | --- | --- | --- | --- | --- | --- | --- | --- | --- | --- | --- | --- | --- | --- | --- |
| NA | 0.9815 | 0.9833 | 0.9620 | 0.9528 | 0.9778 | 0.8313 | 0.9821 | 0.9498 | 0.9720 | 0.9758 | 0.9599 | 0.9816 | 0.9649 | 0.9742 | 0.9606 |
| Neither validated | 0.9677 | 0.9839 | 0.9611 | 0.9085 | 0.9493 | 0.7576 | 0.9631 | 0.9315 | 0.9725 | 0.9524 | 0.9624 | 0.9769 | 0.9572 | 0.9532 | 0.9427 |
| One validates | 0.9874 | 0.9823 | 0.9674 | 0.9686 | 0.9863 | 0.8442 | 0.9892 | 0.9496 | 0.9747 | 0.9848 | 0.9522 | 0.9852 | 0.9709 | 0.9826 | 0.9661 |
| Both validate | 0.9869 | 0.9841 | 0.9535 | 0.9554 | 0.9810 | 0.8887 | 0.9883 | 0.9755 | 0.8733 | 0.9790 | 0.9656 | 0.9832 | 0.9686 | 0.9783 | 0.9615 |

1. Parent SNP validation possibilities: **NA** = results for all animals regardless of parentage validation status; **Neither validated**=neither parent validated, either parent not SNP genotyped or SNP failed; **One validates** = one parent SNP validates other SNP failed or not SNP genotyped; **Both validate** = both parents SNP validate
2. Breed abbreviations defined in table S1

We keep a record of the number of SNP mismatches for all parentage validation checks against the listed parents. By 20/12/2016 775,390 parentage validation checks had been performed for 578,963 genotyped animals. 196,428 animals had parentage validation performed on both listed parents. Of them 6.10% of the listed parents failed with 13 or more SNP misconcordances, 0.02% of the listed parents fell into a ‘grey zone’ with 5-12 SNP misconcordances, and 93.88% of the listed parents validated with 0-4 SNP misconcordances. Of the validated parents 94.40% had 0 SNP misconcordances, 5.30% had 1 SNP misconcordances, 0.26% had 2, 0.03% had 3, and <0.01% had 4.

Analysis of the 46 parentage doubtful ‘grey zone’ animals using all available SNP resulted in 10 animals having a different result when all SNP were used versus the ICBF800 (Table S3). These 10 animals had either7, 8, 9, or 10 SNP misconcordances on the ICBF800 panel. Animals with 5, 6, 11, or 12 SNP mismatches on the ICBF800 panel had the same parentage results for when all SNP were used. If a parentage validation or prediction results in 5-12 SNP mismatches, ICBF now analyses all SNP available and uses a ≤1% misconcordance rate to assign or fail parentage. Considering that those 10 animals mean that the ICBF800 panel has made a parentage error of <0.001% this indicates that it is a very accurate panel to use. By having a 2-step parentage validation process for any animal with 0.5-1.5% misconcordance from ICBF800 parentage SNP, provides ICBF with an extremely accurate parentage validation process while minimizing the computational requirements.

### Mating Validation

In May, 2017 ICBF had ~950,000 animals with a valid genotype, of them 195,299 trios existed with all individuals SNP genotyped and the offspring SNP validates to each parent separately. Mating SNP misconcordances (MSM), where the calf was AB and both sire and dam were homozygous for the same allele, were recorded. Of those 96.964% had 0 MSM, 3.000% had 1-3 MSM, 0.037% had 4-7 MSM, and 0.003% (N=5) had 44 to 74 MSM.

For those 5 animals, from 5 different farms, with >40 MSM, we were able to resolve them using a combination of breed composition, progeny SNP validation, and GRM results along with ICBF’s genotype tracking data. For each farm, the DNA kits were sent for the animal and dam on the same day and they were returned to the lab on the same day. It was determined that the cow and calf’s DNA samples were swapped on the farm. Many of the cows had 2 genotyped progeny where 1 had failed SNP parentage. The failed calf would SNP validate to the other calf based on the ICBF800 panel. One calf was listed as being 11% Shorthorn, 84% Limousine, and 5% unknown; its dam was listed as 22% Shorthorn, 68% Limousine, and 10% unknown; while the sire was listed as 100% Limousine. The breed predictions for the same animals were: calf as 36% Shorthorn, 55% Limousine, 9% unknown; dam as 20% Shorthorn, 75% Limousine, and 5% unknown; the sire was 100% Limousine. Given that the breed prediction for the calf matched the listed breed composition for the dam and vice versa this indicated the calf and dam’s DNA were switched. For the animal’s sire all of them had a listed sire that SNP failed validation and a predicted sire. Most of the dams had a listed sire that SNP failed, or didn’t have a SNP genotype, and all had predicted sires. Essentially the calf’s true sire was predicted as the dam’s sire and vice versa. As the listed and predicted sires were older stock bulls or AI sires the predictions passed our QC checks.

### Parentage SNP Prediction

For any animal who does not have both its sire and dam SNP validate parentage prediction is ran using the ICBF800 panel. This includes animals with a SNP failed listed parent, a SNP ungenotyped listed parent, or no listed parent. Essentially every animal with a valid genotype is checked to see if it could be the parent of the animal. To increase computing speed and keep high accuracy the following procedure is used.

First we create a table that contains the ICBF800 genotypes for all animals, this table is updated daily as new genotypes are received or bad genotypes invalidated. This table contains all animals that have a valid genotype and ≥600 genotypes of the ICBF800 panel each animal is represented only once. The number of SNP mismatches are calculated for each animal in the table, when over 12 mismatches are found the animal has ‘failed’ and is excluded from the analysis. If all 800 SNP are analysed for an animal and there are <12 SNP mismatches then a date of birth and a sex check is performed. If the predicted parent is less than 15 months older than the animal it is excluded, this removes any genotyped progeny of the animal while allowing young uncastrated bulls in the herd to be included. The logic of using 15 months is that the parent was ≥6 months old and 9 months for gestation. While cattle normally reach puberty at 9 to 10 months of age (Rawlings et al., 2008, Gasser et al., 2006), precocious puberty in heifers at 6 months of age has been observed (Wehrman et al., 1996) and there can be variability between an animal’s true and recorded date of birth. The sex check ensures that the animal’s dam is not predicted as its sire and vice versa this also allows an animal’s sire and dam to be predicted together. Any predicted parent with 5-12 mismatches has all available SNP checked after the full table comparison is done.

If more than 1 sire or dam is predicted then all available SNP are used and if only 1 predicted parent has ≤1% SNP misconcordances then that is the predicted parent. If >1 sire and dam have ≤1% SNP misconcordances when all SNP are analysed a manual check is performed. If over 5 sires or dams is predicted then the prediction process stops, without checking all available SNP, and a manual check occurs. In the past >5 sires or dams were predicted because animals were missing an entire genotype class, usually AA or BB.

Predictions are done in batches of 300, so that however many are requested it just peels off the next 300 and runs those. It was found that in the current setup the process was linear, so it was preferred to save off each 300 once done. This way if a prediction process is interrupted for any reason at most only 300 animal’s predictions would be affected.

### MS imputation

While not perfect MS imputation is around 95% accurate (McClure et al., 2014) based upon its use in Ireland. Recently a 91% accuracy for MS imputation was obtained for Slovenia Brown Swiss (Obsteter, 2017) The difference in accuracy may be because ICBF has added over 1,500 more animals to the MS reference population that is publically available in McClure et al. (Mcclure et al., 2013). Animals whose listed parents fail via imputed MS data are directly MS genotyped to confirm or deny the imputed MS result. These animals are then routinely added to the reference MS imputation dataset as they have both SNP and MS genotypes. Imputed MS profiles have also been used as part of the QC pipeline to see if the imputed MS and genotyped MS profiles match. If not then sample is invalidated as the tissue for the SNP profile may not have come from the listed animal. As more animals are genotyped the amount of parentage verification via MS decreases.

### GRM

When parentage cannot be determined via SNP or MS based methods, GRM becomes very useful for identifying close relatives and from that determining a most likely parent. We caution that GRM values can be inflated for inbred animals. GRMs are performed using Golden Helix SVS and the Illumina LD SNP panel. When needed, especially for highly inbred animals, higher SNP density panels are used. Results are used to suggest potential parents of the animal based on the listed pedigree of the identified closely related animals. Suggested parents are not viewed as being validated parents by ICBF, but provide a point of reference for breed societies, farmers, and potential animals to MS parentage check by other groups. Future work will see if GRM values can help identify who a SNP duplicate genotype truly belongs to when parentage validation, offspring validation, sex prediction, and breed composition prediction do not provide a clear answer.

### SNP duplicate and >1 genotype for 1 animal

Both processes use the ICBF800 parentage SNP set and the logic is similar. When an animal’s genotype comes in, if it already has a genotype in the ICBF database the ICBF800 genotypes are compared. If <99% of them match then all available SNP are compared. If <99% pf all SNP do not match then both genotypes are flagged as invalid until resolved.

To identify SNP duplicates the ICBF800 SNP genotypes are converted to a numeric string, for instance 001290021 where 0=BB, 1=AB, 2=AA, 9=missing, and one searches for exact matches of the entire string. Initially, ICBF did use the full string but quickly realized that random missing genotypes would cause some true SNP duplicates to be missed using an exact sting match. Therefore, we broke the 800 SNP string into sixteen 50-SNP non-overlapping blocks and performed exact string matches within each block. If a SNP duplicate was identified for any of the sixteen blocks that pair had all SNP of the ICBF800 compared directly, and separately with all available SNP were compared, missing genotypes excluded. If >99% of the ICBF800 SNP were identical the genotypes were analysed further.

This process worked well, but in February 2016, a SNP duplicate case was identified where random missing SNP caused not even one of the sixteen 50 SNP block to be exactly identical. This lead to using forty 20-SNP blocks to identify potential SNP duplicates. Checking all ~840,000 genotyped animals in March, 2017, the process for identifying potential SNP duplicates took 1 minute to run for 50-SNP blocks compared to 6 minutes for 20-SNP blocks. The 50-SNP block check found 459 potential SNP duplicates and the 20-SNP blocks found 610,172. Once potential SNP duplicates are found all ICBF800 SNP compared SNP by SNP with missing SNP excluded to identify those that are ≥99% identical. This took 13 minutes to run for the 459 potential SNP duplicates and 45 minutes for the 610,172 potential duplicates. Once set up the SNP duplicate check only needs to compare newly submitted genotypes so the time required will greatly decrease. The 20-SNP blocks did find 79 cases of true SNP duplicates that were not identified using the 50-SNP blocks.

For animals that have >1 genotype in the ICBF database the ICBF800 is used to see if ≥99% of the genotypes are identical. If they are not then all available SNP are checked to see if they are ≥99% identical, if not then the genotypes are both flagged as invalid until resolved. For all cases where two genotypes were ≥99% identical for the ICBF800 they were also ≥99% identical for all available SNP. Similarly no case has occurred where >1 genotype per animal was not ≥99% identical for ICBF800 and not also ≥99% for all available SNP.

The genotypes invalidated above are resolved if the results from the parentage validation or offspring validation can clearly identify one genotype as valid and the other as invalid. If not resolved then sex prediction, breed prediction, and date tissue sample submitted information is used. A check on if both animals were on the same farm ever and when tissue samples collected is also performed, if animals have never resided on the same farm then this points to a potential lab caused error. If both animals have resided on the same farm but one was not present when DNA was collected from the other this can be used to resolve whose genotype is valid and whose is invalid.

#### ChrX PAR determination

The IDBv3 has 491 SNP that map to chrX on the UMD3.1 assembly. After filtering for SNP that map to the same location 467 remained. ChrX SNP heterozygosity rates in 38,564 males with 7 chrY SNP called and 568,841 females with 0 chrY SNP were calculated. After filtering for SNP call rate, <0.90, and MAF, <0.01, 388 SNP remained. The PAR region was determined using the average SNP heterozygosity rates in males. The PAR starts between nucleotide 137,112,504 and 137,330,600 where the SNP heterozygosity rate in males jumps from 0.10% to 19.51% for markers IDBV30X00010239 and ARS-BFGL-NGS-90088, respectively (Table S4), in females the average SNP heterozygosity rates for these markers are 22.96% and 34.16%. Based on this PAR start position, the IDBv3 has 287 SNP with unique locations in the chrX nPAR and 101 SNP in the PAR.

The average male heterozygosity rate for the PAR region is 23.92% while the nPAR region is 0.23% compared to 36.95% and 37.06 in females respectively. Figure 1 shows a clear separation of the nPAR heterozygosity rates for the 37,281 males with 7 chrY genotypes and the 568,641 females with 0 chrY genotypes. When we exclude animals with <90% CR for the chrX SNP, 99.95% of the males have <5% PAR SNP heterozygosity rate and 99.87% of the females 99.87% have ≥10%. The average male heterozygosity rate for the PAR region is 23.92% while the nPAR region is 0.23% compared to 36.95% and 37.06 in females respectively (Figure 1)

**Figure 1.**
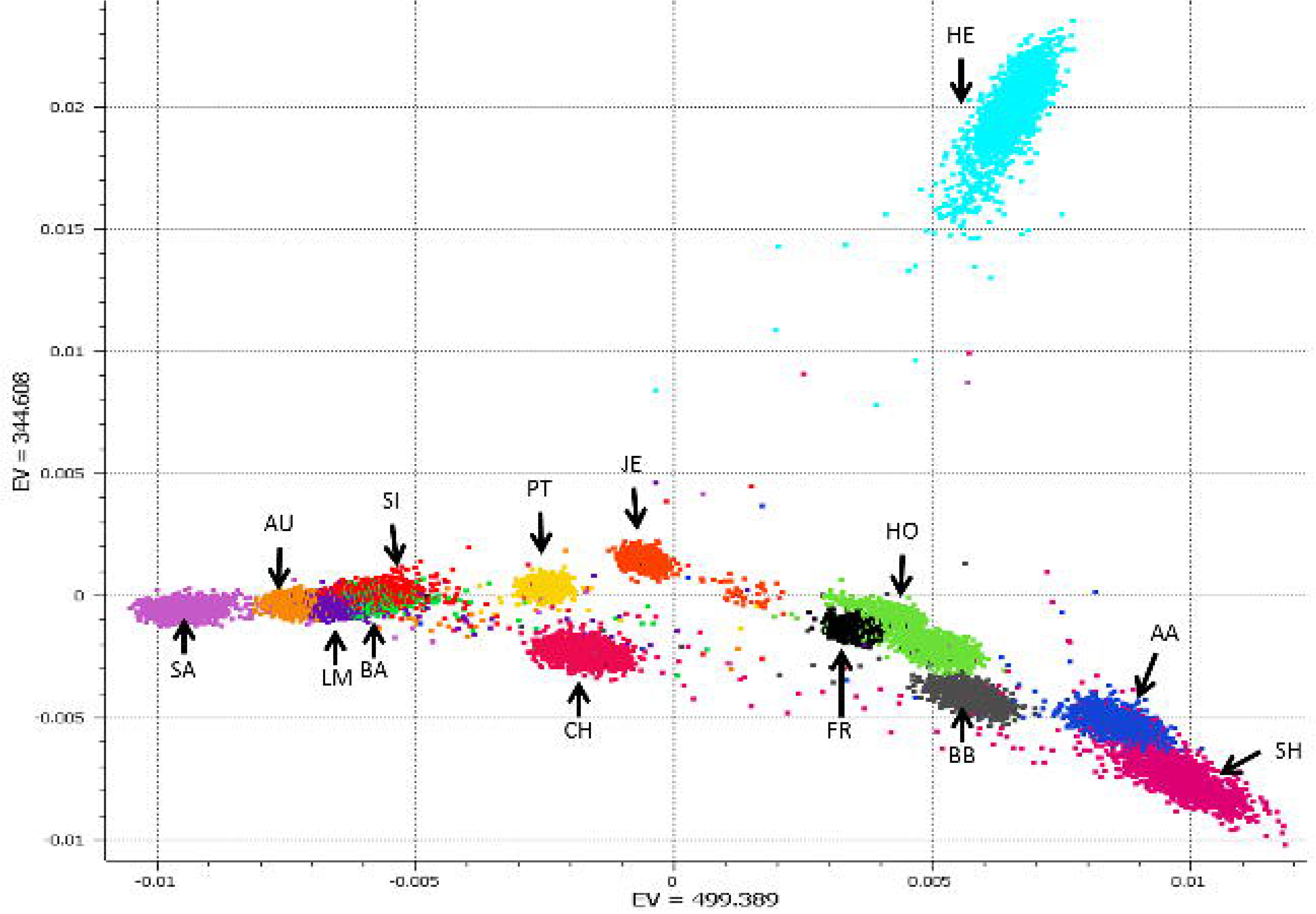
The percent of 37,281 males and 568,641 females at each heterozygosity level for chromosome X SNP that are present pseudosomal region (PAR) and non-pseudosomal regions (nPAR) of chromosome X.

The ChrX nPAR and PAR SNP on the IDBv3 are present on multiple commercial available SNP chips. Across the commercially available SNP chips in the ICBF database (LD, 50k, HD, GGPLDv1, GGPLDv2, GGPLDv3, GGPHDv1, GGPHDv2, IDBv1, IDBv2, and IDBv3) the average number of these nPAR SNP is 225 and PAR SNP is 49 (Table S4).

### Sex Prediction

Sex prediction can be used as an easy QC check to tell if the DNA genotyped truly came from the reported animal, e.g. if a docile cow was sampled instead of a dangerous stock bull. In 612,722 IDBv3 genotyped animals for 7 chrY SNP 568,841 animals had 0 chrY SNP genotypes; 3,109 (1 SNP); 216 (2 SNP); 171 (3 SNP); 138 (4 SNP); 168 (5 SNP); 1515 (6 SNP); and 38,564 (7 SNP). Some research groups have notices chrY SNP genotypes appearing in female Holsteins and this may be caused by an historic event of some part of the chrY transferring to chrX or an autosome (George Wiggins, personal communication, 2016). For the our females with 1 to 4 chrY genotypes 940 have listed dams that have been genotyped on a chip with chrY SNP. Of those 940 animals, which represent a mixture of purebred and crossbred beef and dairy, only 1.81% of their dams also have 0-4 chrY SNP genotypes called, the remaining dams have 0 chrY SNP called (Table S5). Therefore it is very likely that these low levels of chrY genotypes in females is caused by either genotype errors or very low contamination levels.

ICBF predicts an animal’s sex with both nPAR chrY and chrX SNP. Sex prediction is performed initially using 7 nPAR chrY SNP than using 287 nPAR chrX SNP (Table S4). Animals with 6-7 nPAR chrY genotypes or ≤5% heterozygosity rate for nPAR chrX SNP are predicted as male. Females are predicted for those with 0-1 nPAR chrY genotypes or ≥15% heterozygosity rate for nPAR chrX SNP. All others have their sex predicted as ambiguous. Future versions of the IDB will be designed to have more nPAR chrY SNP in hopes to reduce the number of sex ambiguous predictions. Less than 0.001% of animals have an ambiguous sex predicted with our current pipeline.

The current logic used by ICBF for sex prediction is as follows:

1. Predict sex with nPAR chrY SNP

a. Count nPAR chrY genotypes
b. If 0-1 genotypes = female
c. If 6-7 = male
d. If 2-5 = ambiguous sex
2. Predict sex with nPAR chrX SNP

a. Determine heterozygosity rate (# AB/ (#AA + #AB + #BB)) for nPAR SNP
b. If ≤5% het rate = male
c. If ≥15% female
d. If between 5% and 15% = ambiguous sex
3. If sex prediction from step 1 and 2 match report sex predicted
4. If step 1= ambiguous and step 2=male or female (or inverse) then report non-ambiguous predicted sex
5. If step 1 and 2 are both ambiguous = report ambiguous sex predicted

Animals with 0-1 chrY SNP and ≥95% homozygous for all chrX SNP (PAR and nPAR) along with animals with 6-7 chrY SNP and <u>>15</u>% chrX nPAR heterozysitiy rate are flagged for further analysis. These animals could be Turner (X0) or Klinefelter’s yndrome (XXY) animals (Berry et al., 2017).

### Breed prediction

Breed composition prediction is performed for animals whose listed breed composition is comprised of one of 14 reference breeds. Even when >10,000 genotypes are received on a daily basis, ADMIXTURE runs fast enough to be used daily in a production setting. At ICBF breed composition prediction is mainly used as a QC check to tell if the DNA genotyped truly came from the reported animal. As ADMIXTURE provides breed composition percent, future work will look into if this can be applied to help improve genomic breeding value predictions. Correlations of 96.6% have been obtained for of the main breed percent between the ICBF database and 710,000 animals with breed composition predicted (Table 2). Animals that are composed of breeds not in the reference population get predicted as a seemingly random mixture of the reference breed. Up to 24 breed composition prediction was tried with additional breeds having <100 genotyped purebreds (some only 5 animals). Breed prediction for these additional breeds was not accurate (data not shown). PCA analysis of the selected animals from the 14 breeds showed good separation between breeds (Figure 2).

**Figure 2.**
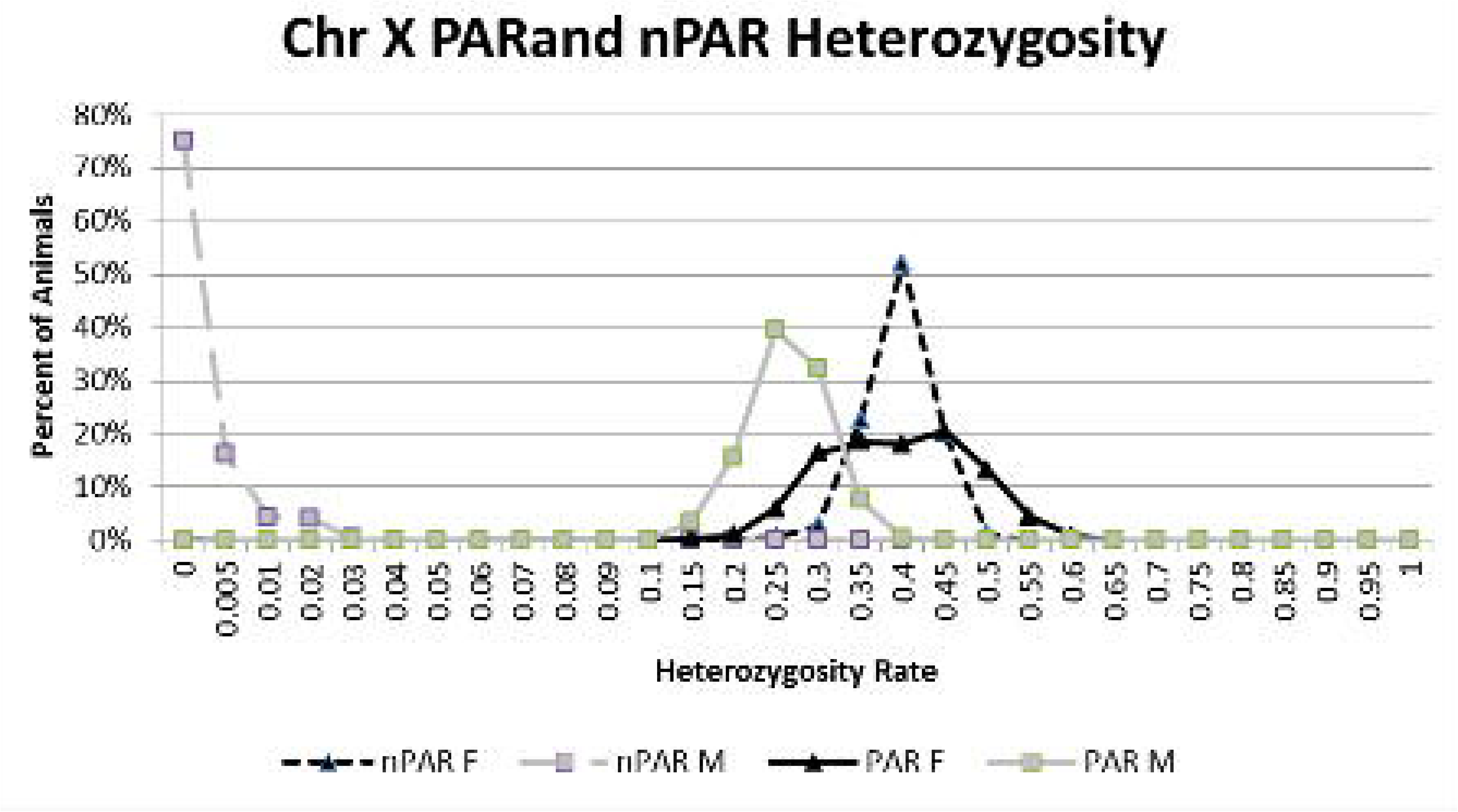
Plot of principle components (PC) 1 and 2 from a PCA analysis of the 22,610 reference animals from 14 breeds. Breed abbreviation listed in Table S1

As shown in Figure 2, most of the breed prediction reference animals fit into close breed specific clusters. There are examples of some animals that are plotted away from their breed cluster. As all reference animals are also ran back through breed composition prediction we can see how their predicted breed compares to the listed breed composition in the ICBF database. If you look at Figure 2 you’ll notice a red dot in the middle of the space between the HE, HO, and JE clusters. According to the ICBF database this animal is 100% Charolais, while according to both the PCA plot and the Admixture results it is predicted to be 50% Charolais and 50% Hereford (the light blue cluster at the top). ICBF will investigate these ‘stray’ PCA results to see if the animal is possible not a true purebred and should be excluded from the reference population. Even with these ‘stray’ animals it is impressive how accurate the breed predictions are, which we believe are because we chose to use a relatively large number of animals per breed.

The accuracy correlation should be looked at with 3 notes: 1) The ICBF database breed composition is based on 32 discrete parts. So a purebred is 32/32 and each part (1/32) is a 3.125% step. ADMIXTURE’s predictions are continuous, so it can predict an animal to be 87.4356% Angus; 2) The reported animal’s breed composition in the database is not perfect; 3) The database calculation assumes that a ½ AA ½ HE breed to a ½ Lim would generate a ¼ AA ¼ HE ¼ HO ¼ LM animal, as we know that is not true.

## Genotype QC Pipeline

The following SNP QC pipeline has been developed at ICBF. While multiple allele formats are available for reporting ICBF typically requests Illumina AB format. Other formats are accepted, but are recoded to AB format before loading. Recoding is based on allele coding information from SNPchiMP (Nicolazzi et al., 2015, Nicolazzi et al., 2014). Overall if a QC step fails then the animal’s genotype is invalidated from the ICBF system. QC is performed at the animal and chip level. The developed pipeline is listed below:

1. SNP call rate: ≥0.90
2. SNP genotypes

A. Are all three AB genotype alleles present (AA, AB, BB) and excluding missing are any other alleles present. If Top or Forward alleles are only A, T, C, G, or missing alleles present
B. Are the AA, AB, BB genotype rates reasonable? If any <20% flag for analysis before using
3. Same ID—same genotypes. If animal is genotyped >1 times then do ≥99% of the called ICBF800 SNP genotypes match
4. SNP duplicate check. Do >1 animals have ≥99% identical genotypes for the ICBF800
5. Sex check. Does predicted sex match the listed sex
6. Breed composition. Does the reported breed and predicted breed composition match
7. Predicted parentage checks:

A. If dam present on farm at conception was predicted stock bull on farm at conception
B. Was predicted dam on farm at birth, three day window allowed
C. Is predicted sire geography located on or close to farm at time of conception
D. If both parents SNP validate, are there >1% ICBF800 SNP misconcordances where both parents are homozygous for the same allele where the offspring is heterozygous
8. Offspring check for validated animals

A. Sires—if ≥5 offspring genotyped and >80% fail then flag sire for manual checking
B. Dam—if <u>>2</u> offspring genotyped and 100% fail then flag dam for manual checking
9. Was tissue sample sent in >45 days after animal’s death

## Discussion and Conclusion

While the ISAG100 and ISAG200 SNP panels do provide a good base for parentage validation via SNP they are not without their limitations, and only using them can result in parentage errors (McClure et al., 2015, Strucken et al., 2015, Strucken et al., 2014, Buchanan, 2016). The number of parentage SNP used by each laboratory, breed society, or national valuation centre will depend on cost and their level of acceptable risk for a parentage error. As the cost of SNP genotyping decreases the value of having a near perfect pedigree could soon outweigh the cost of genotyping an animal with additional SNP. To ensure a higher level of parentage accuracy we recommend that at least 500 SNP are used and more is better. Some might argue that if one restricts parentage to only herd level, less SNP could be used for prediction, but this does not take into account potential errors from fence jumping breeding stock or mis-recorded semen straws. In Ireland, the ICBF800 parentage SNP set have proven very effective for highly accurate parentage validation and prediction, while not being too computationally demanding. Pedigree errors based upon the listed parents in Ireland is runs between 6-8%, is the average rate when all animals regardless of breed, age, or pedigree status are analysed. To date when all QC steps are used, >1 sire or dam has been predicted only due to identical twin animals. While identical twins will have unique methylation patterns (Kaminsky et al., 2009) a method to use this for parentage validation in livestock has not been developed. The ICBF800 is also useful for other processes: such as identification of SNP duplicates, if an animal’s multiple genotypes match, GRM-lite, and parentage prediction as mentioned above. As the ICBF800 were selected based on their performance in a *Bos taurus* population it is unknown how well they would perform in a *Bos indicus* or *Bos taurus* x *indicus* population.

Overall the ISAG800 panel is more accurate than the current international bovine SNP panels; ISAG100 and ISAG200 (Table 3). On its own the ISAG800 panel is not perfect, but by using a 2 step process to further analyse any parentage result with 8-12 SNP misconcordance allows for a highly accurate parentage process with minimal computing requirements. We also recommend not using the SNP we identified having clustering issues for parentage analysis (Table S2). If one wishes to design their own parentage SNP panel we also recommend analysing the SNP’s clustering panel once a large number of animals are genotyped.

**Table 3.** Comparison results of parentage validation test for 300,020 animals between the ICBF800 and smaller SNP panels.

|  | Count<sup>1</sup> | SNP panel used |  |  |  |  |  |  |
| --- | --- | --- | --- | --- | --- | --- | --- | --- |
|  |  | 100<sup>2</sup> | 200 | 300 | 400 | 500 | 600 | 700 |
| Sire | 292462 | 0.249% | 0.022% | 0.006% | 0.003% | 0.002% | 0.001% | 0.001% |
| Dam | 27330 | 0.040% | 0.004% | 0.004% | 0.004% | 0.004% | 0.004% | 0.004% |
| Total<sup>3</sup> | 319792 | 0.231% | 0.021% | 0.006% | 0.003% | 0.002% | 0.001% | 0.001% |

1. Count of SNP parentage checks analysed, by sire, dam or total. 2. Percent of animals that had a different parentage verification result for the 100 SNP panel when compared to the ICBF800 panel at the 1% misconcordance level. 3. 19,774 animals had both sire and dam SNP checked

In the near future national animal registration could occur via SNP genotypes. While an individual’s date of birth can’t currently be determined via its genotype its sex, parents, and breed composition can. SNP genotypes when combined with a robust QC pipeline and a tagging system that collects a tissue sample at the same time a national ID ear tag is applied would allow for unparalleled animal traceability. In theory the animal and its products could be 100% identified if enough genotypes are used which would have applications in animal forensics, theft, and product marketing using a slight modification of the SNP duplication portion of the pipeline.

The ICBF800 works extremely well for parentage and when combined with the rest of our QC pipeline has practically removed the possibility of accidently validating or failing a pedigree incorrectly. While the ICBF800 works across multiple *Bos taurus* breeds, a different set of >500 SNP could eventually be determined to work better for specific or rare breeds. In fact the current set of 800 SNP used by ICBF is no longer the best set of 800 SNP to use based on the MAF criteria we original used (Table S2). This is caused by changing nation-wide MAF as more animals from more breeds are genotyped. In 2014, 69% of the genotyped animals were Holstein; while by March, 2016 Holsteins represented <14% of the genotyped animals. Table 4, shows the major breed component of animals genotyped at ICBF across multiple years. Currently 25 breeds have ≥10 genotyped purebred animals in the ICBF database, while 21 breeds have <10. Even with changing nation-wide MAF the nonISAG200 ICBF800 SNP have an average MAF of 0.484 with a minimum MAF of 0.429 across 852,087 animals on April 5^th^, 2017. When only purebred animals are considered within breeds with >500 animals genotyped (N=15 breeds), the minimum MAF for the nonISAG200 ICBF800 SNP within breed ranges from 0.027 to 0.144 and the average MAF ranges from 0.375 to 0.408. For the ISAG200 SNP the minimum MAF within these purebreds ranges from 0.017 to 0.145 and the average MAF ranges from 0.310 to 0.402 (Summary in Table 5, by SNP in Table S6).

**Table 4.** Percent of major breed represented the total ICBF genotype database by date.

| Breed<sup>1</sup> | 6/2014 | 3/2015 | 3/2016 | 4/2017 |
| --- | --- | --- | --- | --- |
| AA<sup>2</sup> | 4.41 | 7.22 | 9.51 | 9.58 |
| AU | 1 | 0.53 | 0.60 | 0.61 |
| BA | 0.04 | 0.59 | 0.77 | 0.81 |
| BB | 1.01 | 2.75 | 3.63 | 3.63 |
| CH | 9.09 | 19.2 | 19.94 | 21.48 |
| HE | 3.23 | 4.73 | 5.13 | 5.19 |
| HO | 68.02 | 30.08 | 13.89 | 13.63 |
| JE | 0.17 | 0.67 | 0.81 | 0.52 |
| LM | 9.72 | 22.93 | 30.67 | 31.5 |
| MO | 0.14 | 0.05 | 0.16 | 0.17 |
| PI | 0.54 | 0.17 | 0.21 | 0.21 |
| PT | 0.17 | 0.64 | 0.67 | 0.7 |
| SA | 0.04 | 0.92 | 1.70 | 1.72 |
| SH | 0.06 | 1.57 | 1.90 | 1.76 |
| SM | 2.34 | 6.77 | 8.12 | 7.75 |
| Total<sup>3</sup> | 99.98 | 98.82 | 97.7 | 99.26 |

1. The breed represents the animal's major breed component, so an animal that is 75% LM and 25% HO is counted as a LM individual. 2. Breed abbreviations defined in table S1 3. Total values do not add up to 100% as not all breeds are represented

**Table 5.** MAF summary of parentage SNP across 852,087 animals and for 25 breeds

|  |  | ALL | PURE<sup>1</sup> | CROSS<sup>2</sup> | AA<sup>3</sup> | AU | BA | BB | CH | DX | FR | HE | HO | JE | KE | LH | LM | MH | MO | MY | NR | PI | PT | RM | SA | SH | SI | SP | ST |
| --- | --- | --- | --- | --- | --- | --- | --- | --- | --- | --- | --- | --- | --- | --- | --- | --- | --- | --- | --- | --- | --- | --- | --- | --- | --- | --- | --- | --- | --- |
| SNP Panel<sup>4</sup> | Count<sup>5</sup> | 852087 | 146094 | 705993 | 19077 | 2189 | 1354 | 2103 | 34538 | 21 | 710 | 10677 | 16795 | 1315 | 11 | 10 | 39148 | 13 | 204 | 16 | 21 | 595 | 1091 | 59 | 3341 | 2387 | 9900 | 63 | 26 |
| ICBF800 | Max | 0.500 | 0.500 | 0.500 | 0.500 | 0.500 | 0.500 | 0.500 | 0.500 | 0.500 | 0.500 | 0.500 | 0.500 | 0.500 | 0.500 | 0.500 | 0.500 | 0.500 | 0.500 | 0.500 | 0.500 | 0.500 | 0.500 | 0.500 | 0.500 | 0.500 | 0.500 | 0.500 | 0.500 |
|  | Min | 0.267 | 0.255 | 0.269 | 0.040 | 0.053 | 0.091 | 0.058 | 0.073 | 0.000 | 0.082 | 0.017 | 0.082 | 0.017 | 0.000 | 0.000 | 0.051 | 0.000 | 0.069 | 0.000 | 0.000 | 0.114 | 0.132 | 0.000 | 0.027 | 0.050 | 0.035 | 0.024 | 0.038 |
|  | Ave | 0.470 | 0.459 | 0.471 | 0.349 | 0.380 | 0.390 | 0.368 | 0.404 | 0.334 | 0.387 | 0.337 | 0.398 | 0.327 | 0.308 | 0.215 | 0.393 | 0.311 | 0.366 | 0.359 | 0.348 | 0.388 | 0.388 | 0.312 | 0.349 | 0.340 | 0.366 | 0.328 | 0.377 |
| Non ISAG | Max | 0.500 | 0.500 | 0.500 | 0.500 | 0.500 | 0.500 | 0.499 | 0.500 | 0.500 | 0.500 | 0.500 | 0.500 | 0.499 | 0.500 | 0.500 | 0.500 | 0.500 | 0.500 | 0.500 | 0.500 | 0.500 | 0.500 | 0.500 | 0.500 | 0.500 | 0.499 | 0.500 | 0.500 |
|  | Min | 0.428 | 0.345 | 0.435 | 0.040 | 0.109 | 0.091 | 0.105 | 0.144 | 0.000 | 0.083 | 0.028 | 0.124 | 0.034 | 0.000 | 0.000 | 0.051 | 0.000 | 0.069 | 0.000 | 0.000 | 0.130 | 0.132 | 0.000 | 0.027 | 0.072 | 0.094 | 0.024 | 0.058 |
|  | Ave | 0.484 | 0.470 | 0.485 | 0.354 | 0.385 | 0.394 | 0.371 | 0.408 | 0.337 | 0.392 | 0.338 | 0.396 | 0.332 | 0.312 | 0.219 | 0.401 | 0.312 | 0.364 | 0.364 | 0.349 | 0.392 | 0.390 | 0.312 | 0.352 | 0.342 | 0.373 | 0.333 | 0.377 |
| ISAG | Max | 0.499 | 0.500 | 0.498 | 0.499 | 0.499 | 0.500 | 0.500 | 0.498 | 0.500 | 0.499 | 0.497 | 0.498 | 0.500 | 0.500 | 0.500 | 0.499 | 0.500 | 0.500 | 0.500 | 0.500 | 0.499 | 0.499 | 0.500 | 0.500 | 0.499 | 0.500 | 0.500 | 0.500 |
|  | Min | 0.267 | 0.255 | 0.269 | 0.042 | 0.053 | 0.109 | 0.058 | 0.073 | 0.000 | 0.082 | 0.017 | 0.082 | 0.017 | 0.000 | 0.000 | 0.052 | 0.000 | 0.071 | 0.000 | 0.000 | 0.114 | 0.145 | 0.017 | 0.038 | 0.050 | 0.035 | 0.024 | 0.038 |
|  | Ave | 0.427 | 0.422 | 0.428 | 0.334 | 0.363 | 0.374 | 0.356 | 0.389 | 0.327 | 0.374 | 0.334 | 0.402 | 0.310 | 0.296 | 0.201 | 0.370 | 0.308 | 0.371 | 0.340 | 0.345 | 0.376 | 0.381 | 0.313 | 0.339 | 0.332 | 0.346 | 0.315 | 0.376 |

While the SNP part of the QC pipeline results in a ‘black or white’ answer, the Animal QC portion has all parts run and the combined output is used to determine if a genotype should be invalidated. For instance a young animal could pass all the QC checks except sex. This could simply be because the famer recorded its sex wrong by accident. The full Animal QC analysis also provides greater evidence that a genotype truly does not belong to the listed animal if an inquiry ever arises. While sex prediction can be done using only chrX or chrY SNP, we do recommend that both be used to increase sex prediction accuracy and to also identify Klinefelter’s and Turner syndrome animals.

Only after samples have passed the full QC pipeline does ICBF use the data for parentage analysis, genetic disease/trait status, and SNP imputation. The IDBv3 has >200 diagnostic probes for genetic diseases and traits (http://www.icbf.com/?page_id=2170). FImpute (Sargolzaei et al., 2011) is used to impute all animals with a valid genotype to 50k density for genomic breeding value estimation and to IDBv3 density for genetic disease/trait status. By using only valid genotypes and having a highly accurate pedigree via the ICBF800, ICBF is able to help farmers maximize their genetic gain while minimizing their genetic disease risk.

In addition to using ≥500 SNP for parentage validation, we also strongly suggest each laboratory, breed society, or national valuation centre put in place a QC pipeline. The QC pipeline we describe above can be used as an initial foundation for one to build a custom QC pipeline one. One advice we have for any QC pipeline is to include logic checks for any situation one can think of regardless of how ‘rare’ you may think it could occur. It is far easier to deal with a problem genotype early than down the road after pedigrees have been validated, breeding values estimated, and breeding decisions made. The SNP and Animal QC process developed at ICBF has been extremely useful to improve and guarantee the accuracy of the data and any report based on it, from parentage to genomic breeding values. We hope that other can use our QC process to help improve their own systems as they increase their amount of genetic and pedigree data.

## Acknowledgements

We wish to thank all of the Irish farmers that have had their animals genotyped and provided data to ICBF, without them this work would not have been possible.

---

**Figure S1.**
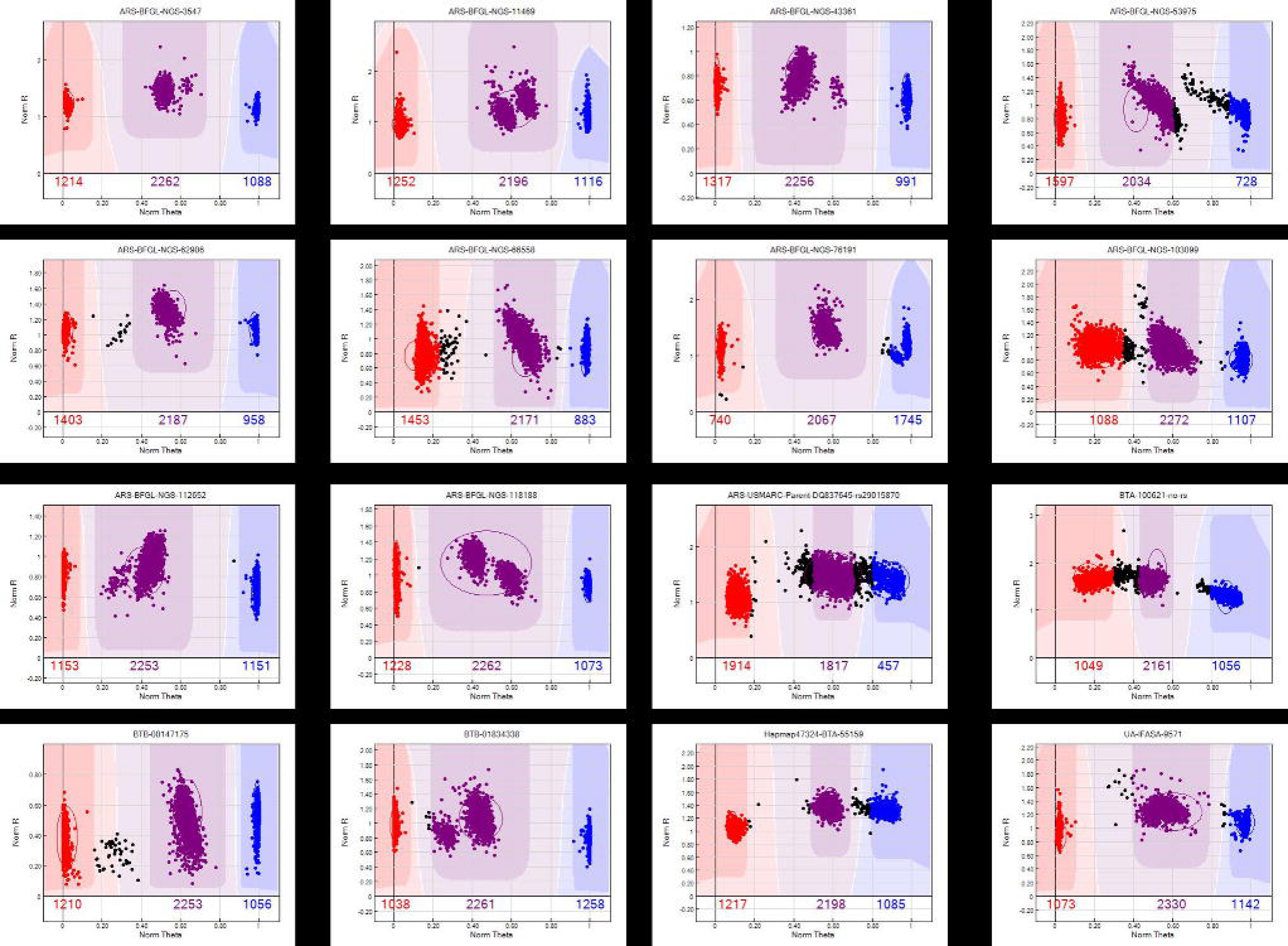
SNP cluster issues for 16 SNP based on Illumina GenomeStudio SNP cluster plots of 4561 animals from multiple breeds. SNP ID (Left to Right and Top to Bottom) are: ARS-BFGL-NGS-3547, ARS-BFGL-NGS-11469, ARS-BFGL-NGS-43361, ARS-BFGL-NGS-53975, ARS-BFGL-NGS-62906, ARS-BFGL-NGS-66558, ARS-BFGL-NGS-76191, ARS-BFGL-NGS-103099, ARS-BFGLNGS-112652, ARS-BFGL-NGS-118188, ARS-USMARC-Parent-DQ837645-rs29015870, BTA-100621-no-rs, BTB-00147175, BTB-01834338, Hapmap47324-BTA-55159, UA-IFASA-9571.

